# High impact journals in ecology cover proportionally more statistically significant findings

**DOI:** 10.1101/311068

**Authors:** Silvia Ceausu, Luís Borda-de-Água, Thomas Merckx, Esther Sossai, Manuel Sapage, Murilo Miranda, Henrique M. Pereira

**Author notes:** corresponding author: S. Ceausu.

## Abstract

Unbiased scientific reporting is crucial for data and research synthesis. Previous studies suggest that statistically significant results are more likely to be published and more likely to be submitted to high impact journals. However, the most recent research on statistical significance in relation to journal impact factors in ecological research was published more than two decades ago or addressed a small subset of the literature. Here, we extract p-values from all articles published in 11 journals in 2012 and 2014 across a wide range of impact factors with six journals sampled in both years. Our results indicate that the proportion of statistically significant results increases with rising impact factor. Such a trend can have important consequences for syntheses of ecological data and it highlights the importance of covering a wide range of impact factors when identifying published studies for data syntheses. This trend can also lead to a biased understanding of the probability of true effects in ecology and conservation. We caution against the possible downplaying of non-significant results by either journals or authors.

## Introduction

Research synthetizing published data in ecology and biological sciences is growing (Newbold et al., 2015) but the validity of its results depends on unbiased reporting of research, including of statistically non-significant results. Incentives in academia that emphasize citations indices and publications in high impact factor (IF) journals may undermine this requirement. For instance, research suggests that statistically non-significant results are less likely to be published in the case of clinical trial studies (2) and are submitted to lower IF journals in ecology (Koricheva, 2003; Suñé, Suñé & Montoro, 2013). Such a bias in publication can result in overestimating statistical significance and effect size in research synthetizing data to estimate a particular effect or phenomenon. Moreover, statistically non-significant results can be highly significant scientifically. Research on the effect of these behaviours on the overall pattern of reported statistical significance in relation to IF has only been conducted on a small subset of the literature (Koricheva, 2003), on a small range of IF (Jennions & Møller, 2003) or has been published more than two decades ago (Csada, James & Espie, 1996). Here, we update the research on the relationship between significance levels and IF in the ecology and conservation literature within a wide range of IF.

## Methods

We examined how the proportion of reported significant results, expressed as p-values, changed with increasing IF. We divided the range of IF into three intervals: low (IF<4), medium (4≤IF<8), and high (IF≥8) to ensure that we cover a broad range of IF. We then randomly selected at least two journals for each IF interval for both 2012 and 2014 from journals listed in the Science Citation Index Expanded (http://mjl.clarivate.com/cgi-bin/jrnlst/jloptions.cgi?PC=D) under four subject categories: biodiversity conservation, biology, ecology, and evolutionary biology. We examined 11 journals (table in S1 Table) (lowest IF 0.36 to highest IF 17.95), six journals for both years, whilst we examined three for 2014 and two for 2012 alone. We collected all p-values reported in all articles using Examine32 Text Search from Aquila Software. We extracted exact p-values and inexact p-values (e.g. p<0.05). All the inexact p-values were reassigned to six intervals in order to harmonize all the reported values and calculate the proportion of significant results (Table A in S1 Text). Due to the ubiquitous use of 0.05 as the alpha level in ecology, we considered p-values below this level as significant. We calculated the proportion of significant results for exact p-values and all (i.e. exact and inexact) p-values reported in each journal, for each year. We fitted generalized linear mixed effects models (binomial distribution) to the proportion of significant results with IF as fixed effect and journal as random effect. We used R (version 3.4.0) and the “lme4” and “arm” packages for model fitting and back-transformation of model coefficients. We tested through a paired t-test the effect of year on the six journals that we examined for both years. All data collected for this study are available at: https://figshare.com/projects/High_impact_journals_cover_proportionally_more_statistically_significant_findings/28284. The R code of the data analysis is available at: https://github.com/SilviaCeausu/ImpactFactorsAndPvalues/blob/master/ImpactFactorPvaluecodeGitHub2.R.

## Results and discussion

The percentage of significant results–out of all published results within a journal– increased with increasing IF (Figure 1). For exact p-values, the percentage of significant results increased on average by 0.7% for each additional IF unit [95% CI: 0.3 - 1.2%]. For all p-values (i.e. exact and inexact), the percentage increased by 1.2% for each additional IF unit [95% CI: 0.9 - 1.5%] (Table S2). In 2012, the percentage of significant results was higher than in 2014 (+3.3%, 95% CI: 0.2 – 6.4%) for the journals examined in both years.

**Figure 1.**
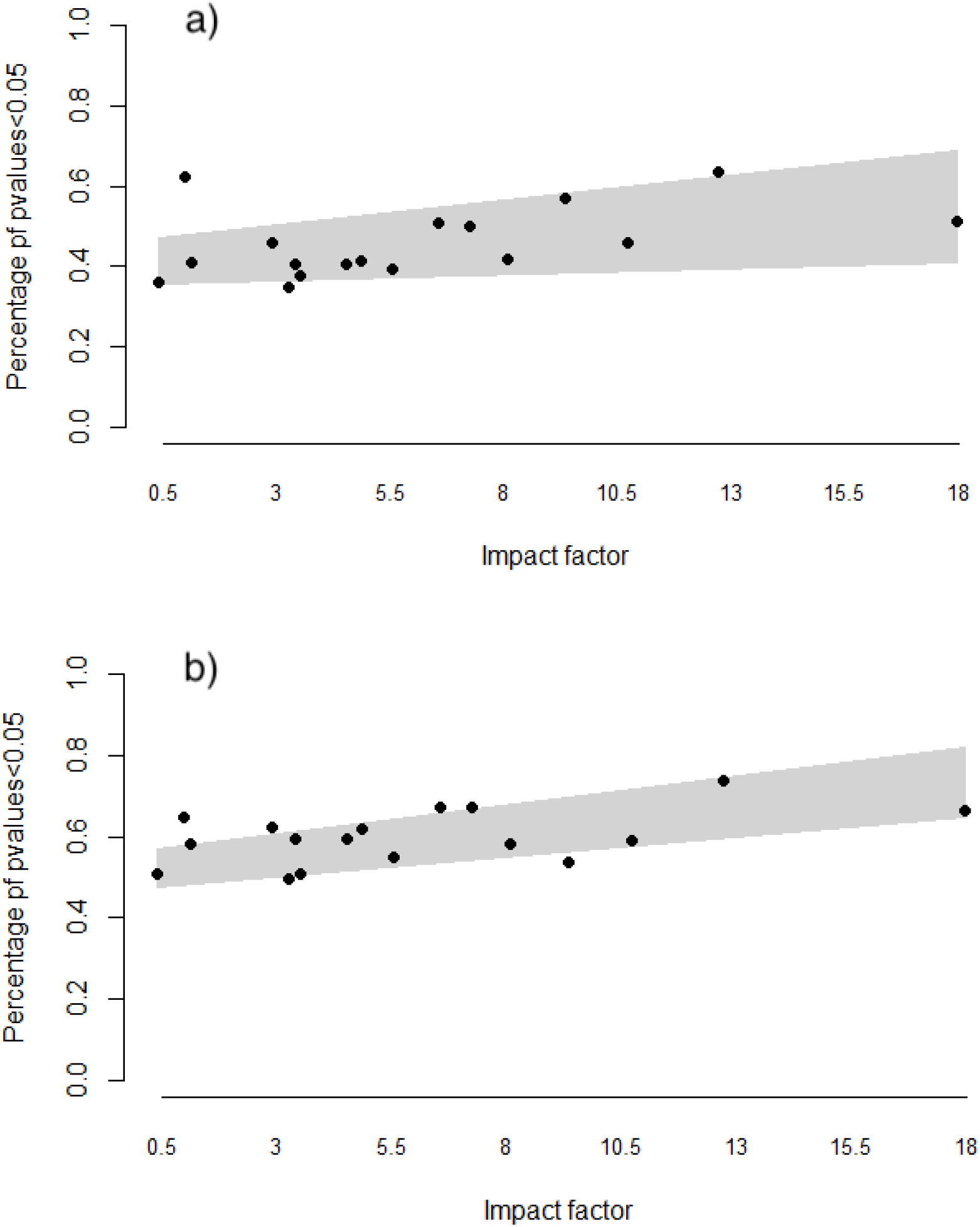
Proportion of a) exact p-values and b) all p-values below 0.05 reported across impact factors (IF). The grey areas delimitate the 95% confidence intervals. The journals considered and their impact factors are presented in S1 Table.

Our result concurs with trends noticed in medical research (Jannot et al., 2013) but they contradict results reported for behavioural ecology by (Jennions & Møller, 2003). The latter study concluded that neither p-values nor statistical power varied significantly with IF but the analysis was conducted on a much narrower range of IF (ca. 1 - 5) than our study.

Our result can arise if statistical significance influences submission or editorial decisions. Analysing the output of doctoral dissertations in ecology, Koricheva (Koricheva, 2003) found that the proportion of non-significant results in a study was negatively associated with IF, although the rejection rates for non-significant results were not higher for higher IF journals. In an article that examined clinical trials, Suñé et al (Suñé, Suñé & Montoro, 2013) found that non-significant studies are less likely to be published and, if published, more time passes between conducting the research and publication. However, the IF of the publishing journal was not different for significant versus non-significant results (Suñé, Suñé & Montoro, 2013). These studies suggest that authors invest less effort into the publication of their non-significant results and submit them to lower IF journals.

Our outcome can also be an effect of higher impact journals selecting studies with higher sample or effect sizes, or requesting stricter statistical reporting and shorter articles. In studies of the relationships between IF, and sample and effects sizes results are mixed. For meta-analyses in ecology Lortie et al (Lortie et al., 2013) did not find a relationship between IF and effect size. Analyzing studies collected for four meta-analyses, Murtaugh found a positive correlation between effect strength and IF in two of the four datasets (Murtaugh, 2002), and Barto and Rilig (Barto & Rillig, 2012) found that high IF journal published the strongest effects, although in the absence of correlations with data quality. Regarding statistical reporting, Tressoldi et al (Tressoldi et al., 2013) suggest that higher IF journal do not necessarily display better standards. Our data also show no indication that higher IF journals publish more precise p-values then lower IF journals (figure in S1 Figure). The heterogeneity of article length requirements across journals did not allow us to test whether article length requirements play a role in our result (table in S1 Table). Moreover, we did not analyse supplementary materials, which might include additional non-significant results considered secondary by authors.

The large confidence intervals in our results suggest that other factors also have an influence on publication. For example, the difference in percentage of significant results between years suggests changes in the prominence of different research topics. However, we cannot exclude an undervaluation of non-significant results, either by authors or by journals. This pattern may make significant results more visible if they are published in higher IF journals than non-significant findings, and may create an inaccurate perception of the probability of true effects in ecology. This could lead to wasted efforts on approaches or interventions that could in reality be far less effective than we assume (Meli et al., 2017). Publication biases could also negatively impact our understanding of biodiversity change and its drivers if a higher proportion of non-significant results remain unpublished compared with significant ones, especially at a time when growing synthesis efforts are trying to shed light on important biodiversity and ecology questions (Vellend et al., 2013; Dornelas et al., 2014; Newbold et al., 2015). Moreover, statistically non-significant results can give rise to new theories or amendments to old ones, as it is the case of the emerging debate on the importance of isolation in fragmented landscapes (Collinge, 2000; Fahrig, 2013). Therefore, we advise careful consideration of submission and publication decisions to ensure solid foundations for our scientific understanding.

## Acknowledgments

MS is funded by a PhD grant from Fundacão para a Ciência e a Tecnologia (FCT), Portugal (ref. PD/BD/128349/2017).

## Supporting information

**Data collected for this study are available at:** https://figshare.com/projects/High_impact_journals_cover_proportionally_more_statistically_significant_findings/28284.

**R code of the data analysis is available at:** https://github.com/SilviaCeausu/ImpactFactorsAndPvalues/blob/master/ImpactFactorPvaluecodeGitHub2.R.

**S1 File. Data collection protocol (separate document)**

**S1 Table.**
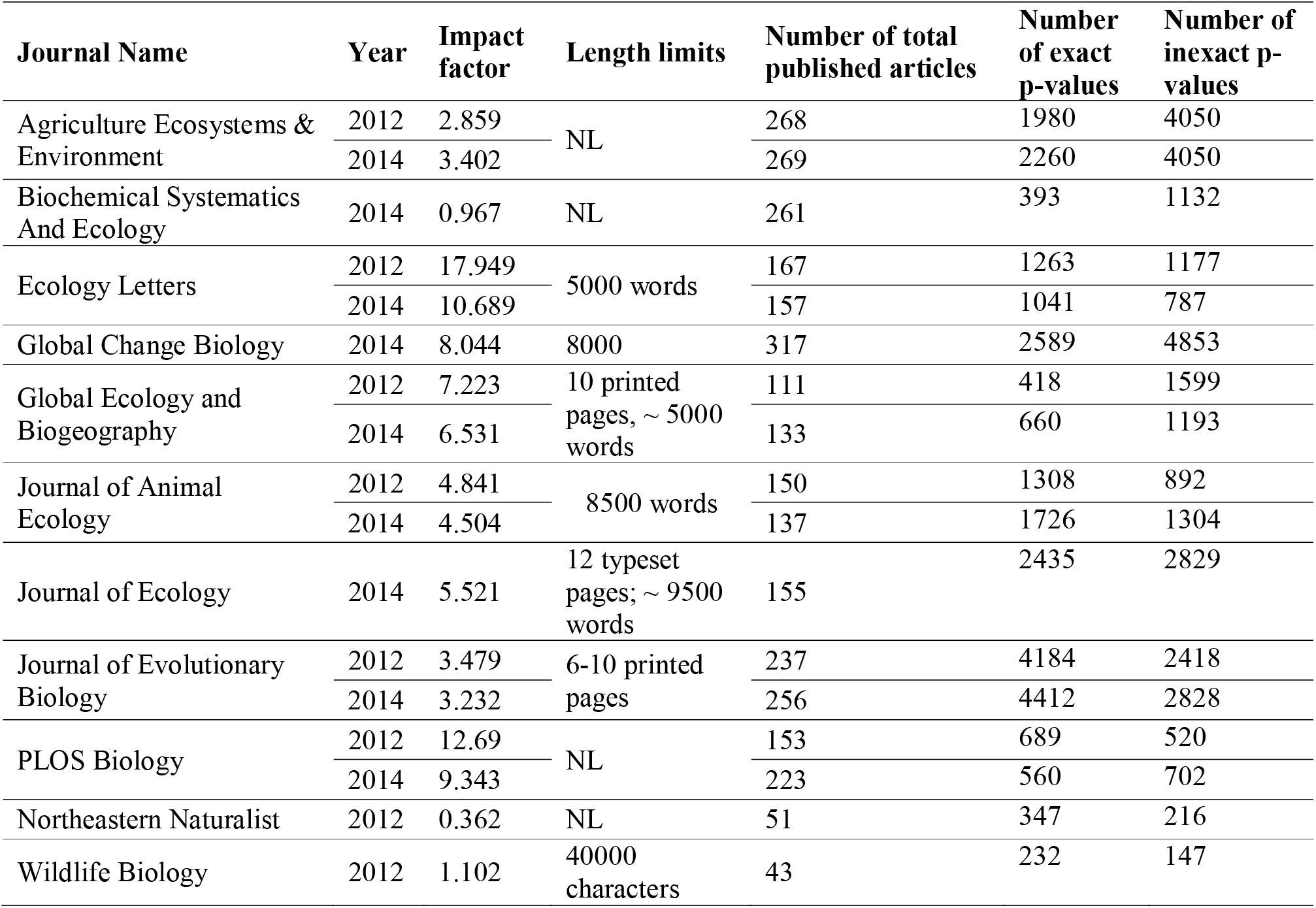
Information regarding the journals included in the analysis. Journal title, publication year, the impact factor in the respective year, length limits for the main article type of the journal, total number of articles published by each journal during the respective year, total number of exact and inexact p-values identified. NL - no length limit specified in the author guidelines.

| Journal Name | Year | Impact factor | Length limits | Number of total published articles | Number of exact p-values | Number of inexact p-values |
| --- | --- | --- | --- | --- | --- | --- |
| Agriculture Ecosystems & Environment | 2012 | 2.859 | NL | 268 | 1980 | 4050 |
|  | 2014 | 3.402 |  | 269 | 2260 | 4050 |
| Biochemical Systematics And Ecology | 2014 | 0.967 | NL | 261 | 393 | 1132 |
| Ecology Letters | 2012 | 17.949 | 5000 words | 167 | 1263 | 1177 |
|  | 2014 | 10.689 |  | 157 | 1041 | 787 |
| Global Change Biology | 2014 | 8.044 | 8000 | 317 | 2589 | 4853 |
| Global Ecology and Biogeography | 2012 | 7.223 | 10 printed pages, ~ 5000 words | 111 | 418 | 1599 |
|  | 2014 | 6.531 |  | 133 | 660 | 1193 |
| Journal of Animal Ecology | 2012 | 4.841 | 8500 words | 150 | 1308 | 892 |
|  | 2014 | 4.504 |  | 137 | 1726 | 1304 |
| Journal of Ecology | 2014 | 5.521 | 12 typeset pages; ~ 9500 words | 155 | 2435 | 2829 |
| Journal of Evolutionary Biology | 2012 | 3.479 | 6-10 printed pages | 237 | 4184 | 2418 |
|  | 2014 | 3.232 |  | 256 | 4412 | 2828 |
| PLOS Biology | 2012 | 12.69 | NL | 153 | 689 | 520 |
|  | 2014 | 9.343 |  | 223 | 560 | 702 |
| Northeastern Naturalist | 2012 | 0.362 | NL | 51 | 347 | 216 |
| Wildlife Biology | 2012 | 1.102 | 40000 characters | 43 | 232 | 147 |

**S2 Table.**
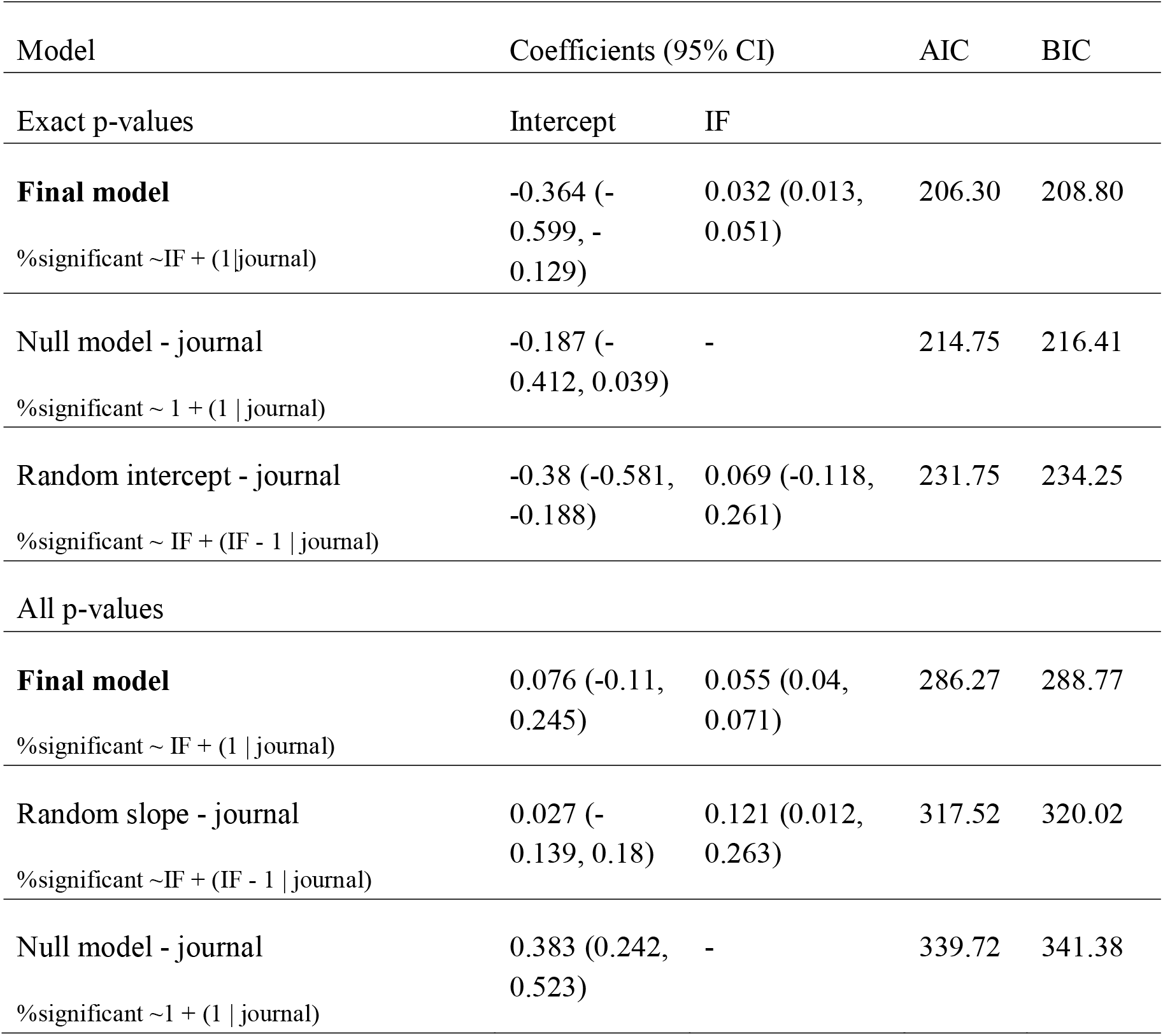
Model coefficients and selection. We tested the model used for Figure1 (main text) against alternative random effects structures and null models. A model with both random slope and intercept was not possible due to the low number of data points. We compared the models according to theoretic information criteria: Akaikes’ Information Criterion (AIC) and Bayesian Information Criterion (BIC). We used the same models for both exact p-values and all p-values. We rounded up all values to three decimal places.

**S1 Figure.**
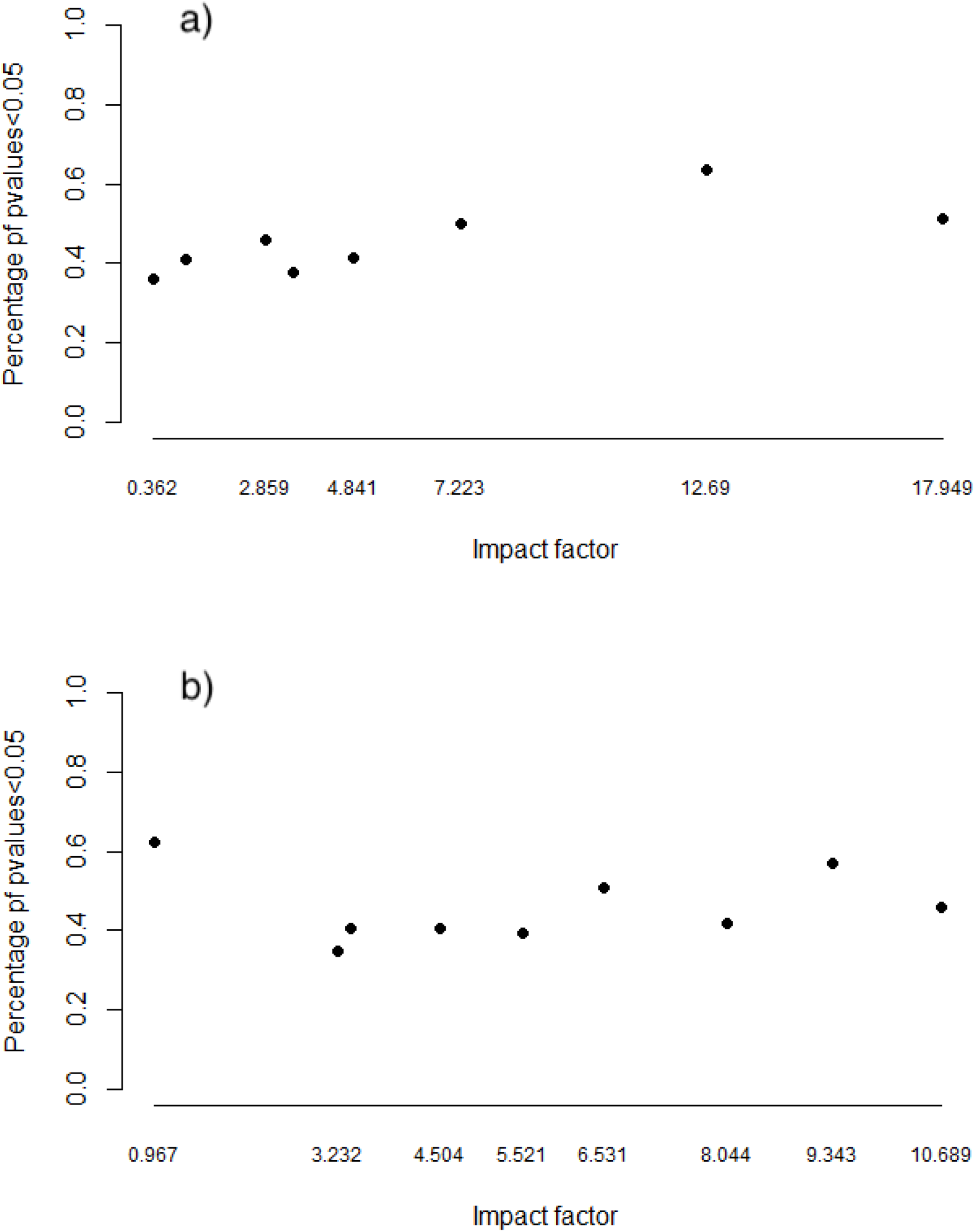
Proportion of exact p-values below 0.05 reported in a) 2012 and b) 2014 across the range of impact factors (IF).

**S2 Figure.**
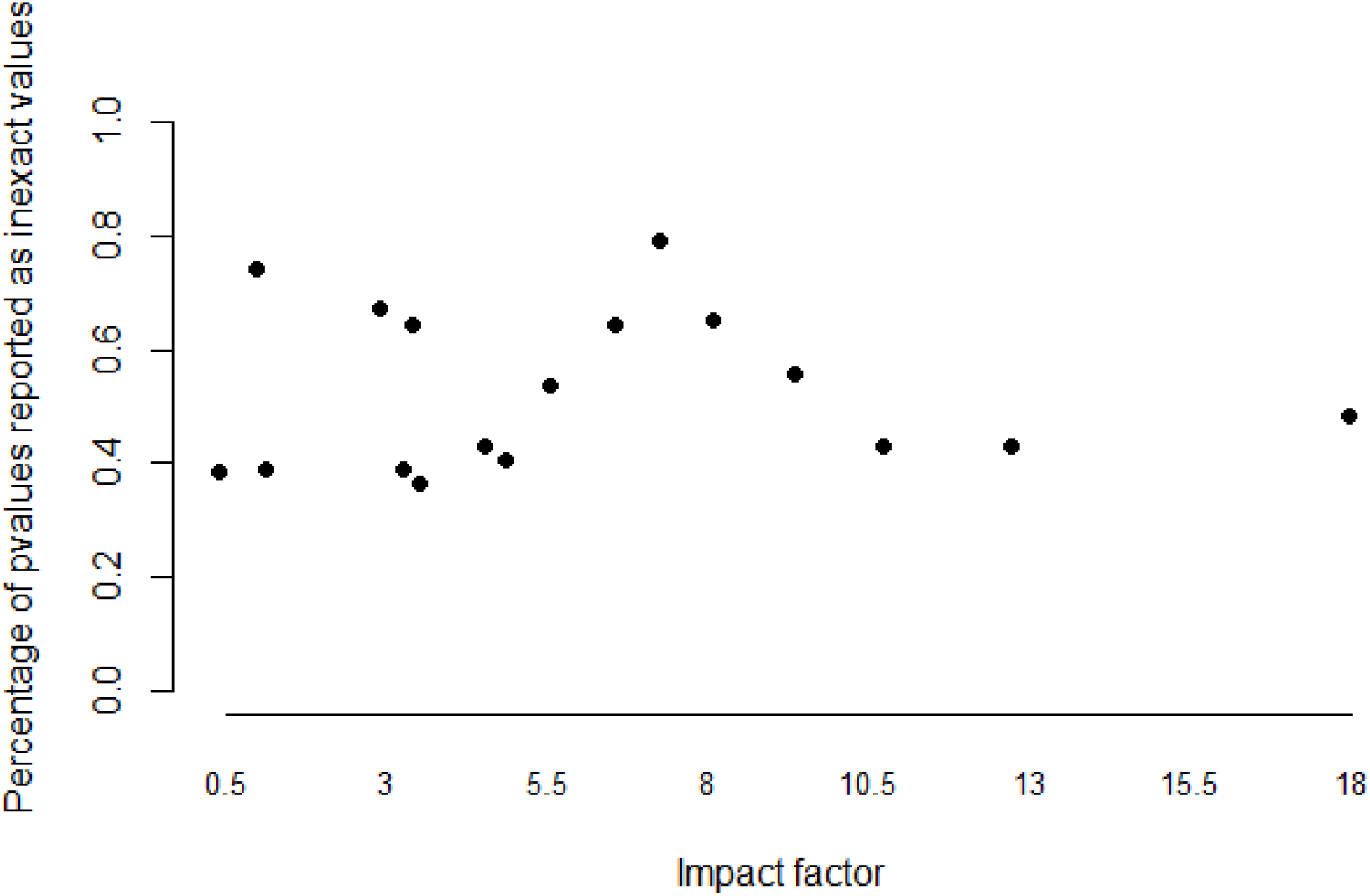
Proportion of inexact p-values out of the total reported values across the range of impact factors (IF).

